# Altered Brain Energy Metabolism in the APPPS1 Alzheimer’s Model during anesthesia: Integration of Experimental Data and In Silico Modeling

**DOI:** 10.1101/2025.06.26.661690

**Authors:** Jonas Schunack, Georg Riepe, Frank L. Heppner, Kathrin Textoris-Taube, Iwona Wallach, Michael Mülleder, Agustin Liotta, Marina Jendrach, Nikolaus Berndt

**Author notes:** Corresponding author Nikolaus Berndt German Institute of Human Nutrition Potsdam-Rehbruecke (DIfE) Department of Molecular Toxicology Nuthetal, Germany. These authors contributed equally to this work (Marina Jendrach, Nikolaus Berndt).

## Abstract

**Background:** Synaptic transmission and network activity rely on high ATP turnover. Impairments in cerebral energy metabolism are increasingly recognized as central in aging and Alzheimer’s disease (AD) pathogenesis. Elderly patients and patients with AD are also at elevated risk for perioperative neurological complications, including post-operative delirium and further cognitive deterioration. However, the interaction between metabolic vulnerability and anesthetic exposure remains incompletely understood.

**Methods:** We investigated cortical metabolic responses and potassium homeostasis in acute brain slices from wild-type (WT) and AD-like APPPS1 transgenic mice, which were either exposed to isoflurane or left untreated. Glia cells were assessed by staining microglia and astrocytes. Measurements of the cerebral metabolic rate of oxygen (CMRO₂), extracellular potassium dynamics, and proteomic profiling were integrated with computational modeling to assess oxidative metabolism and anesthetic effects under different conditions.

**Results:** APPPS1 mice exhibited reduced CMRO₂ and attenuated neuronal activity compared to age-matched WT controls, showing sex-specific differences. Proteomic analysis revealed the downregulation of key mitochondrial and glycolytic enzymes, indicating an impaired ATP-generating capacity. Exposure to isoflurane further suppressed CMRO₂, with a more pronounced effect in the APPPS1 brain tissue, while glia cells exhibited no acute changes. Additionally, isoflurane exacerbated deficits in extracellular potassium ([K⁺]ₒ) clearance, highlighting impaired ion homeostasis under anesthetic challenge.

**Conclusions:** Our findings demonstrate that AD-like pathology in APPPS1 mice is associated with a significant decline in oxidative metabolism and ATP availability. These deficits are exacerbated by anesthetic exposure, contributing to impaired potassium regulation. This suggests that diminished metabolic flexibility may underlie increased anesthetic vulnerability and postoperative complications in AD.

## Introduction

Anesthesia is an essential prerequisite for countless surgical and diagnostic interventions, with more than 300 million anesthetic procedures performed globally each year. While modern anesthetic protocols are generally safe, particularly in younger and healthy individuals, the situation is markedly different in elderly populations and patients with underlying neurological conditions. Among these, postoperative delirium (POD) stands out as a prevalent and serious complication, affecting 10–20% of elderly patients undergoing anesthesia. POD typically manifests within the first few days postoperatively and is characterized by acute disturbances in cognition, attention, and awareness. Though transient, it significantly increases the risk of long-term cognitive decline and disability, particularly in vulnerable individuals [1].

Despite the frequency and severity of POD, its underlying molecular mechanisms remain incompletely understood. Several factors have been proposed, including neuroinflammation, impaired mitochondrial function, and disrupted ion homeostasis [2]. Mitochondria, the primary source of neuronal ATP, are highly susceptible to stress. Impaired mitochondrial function results in decreased ATP availability and increased oxidative stress—both of which can undermine neuronal integrity and synaptic transmission [3]. Glial cells, including astrocytes and microglia, are similarly dependent on ATP not only for basic maintenance but also for active roles in modulating synaptic function [4] and driving inflammatory responses [5].

One of the strongest predisposing factors for POD is a diagnosis of Alzheimer’s disease (AD) [6, 7]. Patients with AD exhibit marked mitochondrial dysfunction, altered energy metabolism, and defective mitochondrial quality control [8, 9]. However, mechanistic studies on the interaction between anesthesia and AD pathology are complicated by the fact that patients often require surgery due to comorbid conditions, introducing systemic confounds. Moreover, *in vivo* models for POD are limited in their ability to isolate central nervous system changes from peripheral influences [10, 11].

To address this, we adapted an acute brain slice model for use in the context of AD-like pathology. This *ex vivo* approach enables the investigation of intrinsic metabolic responses under tightly controlled experimental conditions, preserving local cellular architecture and neural circuitry while avoiding systemic variability [12].

We used brain slices from 11-month-old APPPS1 transgenic mice, a well-established genetic model of AD that presents with both amyloid-beta (Aβ) deposition and progressive neuroinflammation by this age [13, 14]. These slices were exposed to clinically relevant concentrations of isoflurane (1% for light anesthesia and 3% for deep anesthesia), and changes in cerebral metabolic rate of oxygen (CMRO₂), potassium homeostasis, and glial activation were assessed.

Given that sex is a critical biological variable in both AD and anesthetic outcomes [15]—with women exhibiting faster AD progression [16] and men showing higher POD incidence [17, 18]—we performed all analyses in sex-stratified groups. Furthermore, to understand the biochemical basis of our physiological findings, we performed quantitative proteomic profiling and integrated these data into a kinetically resolved model of neuronal energy metabolism [19].

By integrating these modalities, we aimed to investigate how isoflurane anesthesia affects oxidative metabolism, ion regulation, and glial responses in the context of AD-like pathology. Understanding how anesthetics interact with the bioenergetic landscape of the Alzheimer’s brain may help elucidate the mechanisms underlying POD and cognitive decline, and ultimately support the development of strategies to reduce perioperative risk in patients with neurodegeneration.

## Materials and Methods

### Animals

Experimental protocols were approved by German animal welfare authorities (Landesamt für Gesundheit und Soziales, Berlin, T-CH 0039/21). This study was conducted following the ARRIVE 2.0 guidelines, Helsinki declaration, and Charité animal welfare guidelines. Prior to *in vitro* experiments, the animals had a minimum of seven days for acclimation to our animal shed, where they were held in groups of at least two and with food ad libitum and a 12-hour light cycle.

### Slice preparation and maintenance

For *in vitro* experiments, hippocampal slices from male and female 11-month-old wild-type (WT) and transgenic APPPS1 mice [13] were prepared as previously described for rats [12].

Artificial cerebrospinal fluid (aCSF) contained (in mM): 129 NaCl, 21 NaHCO_3_, 10 glucose, 3 KCl, 1.25 NaH2PO4, 1.6 CaCl2, and 1.8 MgCl2. pH was 7.35-7.45 and osmolarity was 295-305 mosmol/L. In brief, animals were placed in an anesthesia chamber and anesthetized using isoflurane (3%) in 100% oxygen to avoid hypoxia (flow 2 L/min) previous to decapitation. Brains were removed and horizontally sectioned into 400 μm-thick slices using a LeicaVT 1200s vibratome (Leica, Wetzlar, Germany) and promptly transferred to a resting interface chamber. Slices were then perfused with carbogenated aCSF (∼36 ± 0.5 °C) and continuously aerated with humidified carbogen (95% O_2_, 5% CO_2_). Brain slices were randomized during allocation in the recording chamber. Experimental procedures commenced after a two-hour stabilization period. At the end of the preparation procedures, 2 slices per animal were separately snap-frozen and stored at -80 °C for proteomic analysis.

### Electrophysiology, ptiO2 recordings

As circuitry and cell distribution are well known, the entorhinal cortex (EC) was chosen to perform electrophysiological recordings. Simultaneously, field potentials, extracellular potassium concentrations ([K^+^]_o_), and partial tissue oxygen pressure (p_ti_O_2_) measurements were performed. Tissue oxygen was measured using Clark-style oxygen (tip diameter: 10 μm; Unisense, Aarhus, Denmark) and [K^+^]_o_ was assessed with double-barreled ion-sensitive microelectrodes constructed and calibrated as reported by Angamo et al. [20]. We used the Potassium Ionophore I 60031 (Fluka, Buchs, Switzerland) accordingly. To create p_ti_O_2_ depth profiles, the oxygen microelectrode was mounted on a mechanical micromanipulator (Narishige, Japan) and moved vertically in 20-μm steps through the brain slice until p_ti_O_2_ reached its minimum as previously established by our group [21].

Activity-dependent changes in pO_2_ and [K^+^]_o_ were elicited by 2-s long 20 Hz tetani (single pulse 100 μs duration, interval 50 ms, 40 pulses) with a bipolar stimulation electrode. Pulse generation was performed with Master 8 (A.M.P.I., Jerusalem, Israel). Tetani were repeated three times to measure activity-dependent changes at 40 µm, 80 µm, and at the core of the slice. These measurements were repeated in four situations: under control conditions, followed by exposure to 1% isoflurane, then 3% isoflurane, and finally after a washout. For all consecutive measurements, a 20-minute (±5 min) baseline stabilization period was maintained between each condition to ensure comparable resting conditions and stable measurement signals.

### Isoflurane application

Using a calibrated vaporizer (Dräger, Germany), Isoflurane was applied in the interface chamber at a gas flow of 1 L/min. Concentration control was performed using a Vamos® mobile isoflurane monitor (Dräger, Germany). Taking into consideration the water/gas partition coefficient for 37 °C of 0.54, the application of 1% and 3% lead to 0.24 and 0.72mM isoflurane in the aCSF [22]. Importantly, 1% and 3% isoflurane are clinically relevant concentrations for light and deep anesthesia, respectively.

### Histology

After isoflurane treatment, 400μm brain sections were permeabilized in PBS/2% Triton-X-100 for 4 h and blocked by incubation at RT for 2 h in PBS/10% goat serum/2% Triton-x-100/2% bovine serum albumin. Primary antibodies were used in a 1:300 dilution in PBS/5% goat serum/0.3% TX) at 4 °C. Primary antibodies were Iba1 (019-19741, Wako), TREM2 (AF1729- SP, R&D Systems), and GFAP (Z0334, Dako). The next day, slices were washed and the corresponding secondary antibodies were added in a 1:300 dilution in PBS/5% goat serum/ 0.3% TX) for 2 h at RT: 568 anti-rabbit (A11011, Invitrogen), 647 anti-sheep (ab150179, abcam), and 488 anti-rabbit (A11008, Invitrogen). Nuclei were stained with DAPI 1:1000 for 1 min (10236276001, Roche) and afterward, slices were mounted. Stainings were imaged using a Leica TCS SP5 confocal laser scanning microscope controlled by LAS AF scan software (Leica Microsystems, Wetzlar, Germany) using a 20x objective. GFAP- and IBA1-covered area and TREM2 intensity were analyzed using ImageJ.

### Data Acquisition and Data Analysis

Analog signals were digitalized using the Power CED1401 and Spike2 software (both from Cambridge Electronic Design, Cambridge, UK). Data analysis and statistical evaluations were conducted with Spike2, Excel (Microsoft, Seattle, USA), MATLAB (MathWorks Inc., Natick, USA), and R Statistics (R Core Team, Vienna, Austria). R and MATLAB scripts automated blinded raw data processing and minimized the potential risk of human error and observer bias, thereby increasing standardization. For CMRO₂, the median values along with the 25th and 75th percentiles (in brackets) are presented in the results. Data is visualized using box plots, displaying the median, mean, and interquartile range (25th and 75th percentile). The absolute [K^+^]_o_ was calculated using a modified Nernst equation and calculations regarding the potassium transients were measured using automated MATLAB scripts. We estimated [K^+^]_o_ clearance by measuring the time of [K^+^]_o_ decay after stimulation during two distinct periods: an early decay time (half decay from maximal amplitude or T50) and a late decay time (90% decay from maximal amplitude or T90) (see also Reiffurth et al. [23]). For statistical inference, we tested for normal distribution and residual normality, followed by repeated-measures ANOVAs (including appropriate error terms to account for within-sample dependencies), Student’s t-tests, and linear mixed-effects models with random intercepts per sample. P values for fixed effects in mixed models were calculated using the Satterthwaite approximation as implemented in the lmerTest package. P values were adjusted using the Bonferroni correction when multiple comparisons were performed. All results were analyzed with respect to the animal’s gender. “N” refers to the number of animals and “n” to the number of slices used for each experimental condition. A significance threshold was set at p <0.05.

### Calculation of Cerebral Metabolic Rate of Oxygen

The cerebral metabolic rate of oxygen (CMRO₂) was calculated based on ptiO₂ depth profiles, following a previously established method [21]. Briefly, we utilized a reaction-diffusion model that accounts for both the diffusive transport of oxygen and its consumption within the tissue slice. To achieve this, the slices were segmented into layers of equal thickness (1 µm each). The diffusion of oxygen between these layers was described using Fick’s Law, assuming a diffusion constant of 1.6 × 10³ µm²/s. Oxygen consumption within each layer followed Michaelis-Menten kinetics, with a Km-value of 3 mmHg [24]. The CMRO₂ was assumed to be homogeneous across the tissue slice and was treated as a variable parameter optimized to achieve the best fit with the experimental data. For the boundary conditions, the ptiO₂ concentration at the slice surface was constrained to the supplied oxygen level, whereas at the ptiO₂ minimum, diffusive oxygen transport was set to zero. Consequently, surface ptiO₂ concentrations were measured at 682 mmHg under control and washout conditions, while reduced values of 675 mmHg and 662 mmHg were observed for 1% and 3% isoflurane, respectively, due to the displacement of oxygen in the supplied gas mixture.

### Proteomics Sample Preparation

For proteomics analysis, ∼10-50 mg of mouse brain tissue was weighed into Lysing Matrix D Tubes (MPBio 116913100) and the volume was made up to 300 µL of RIPA lysis buffer (Thermo 89900) with 1.25x protease inhibitor (Merck 11873580001). Samples were homogenized with three repeats in MP FastPrep instruments, settings 6 m/s, 3x30 sec.

Debris was collected (13000 rpm for 5 min), the protein extractions were analyzed with quantitative BCA Assay (Pierce Protein Assay Kit, 23225), and 25 µg of protein in 50 µL were transferred to a plate. Digestion with Benzonase® HC nuclease (NEB), 40 units per sample incubation was performed at 37 °C for 30 min before SP3 protein preparation. Lysates were processed on a Biomek i7 workstation using the SP3 protocol as previously described with one-step reduction and alkylation [25]. Briefly, 16.6 μL of reduction and alkylation buffer (40 mM TCEP, 160 mM CAA, 200 mM ABC, 4% SDS) was added, and samples were incubated at 95 °C for 5 min and cooled to RT. To bind the proteins, 250 μg of paramagnetic beads (1:1 ratio hydrophilic/hydrophobic) were added and the proteins were precipitated by adding 50% ACN. Samples were washed twice with 80% EtOH and once with 100% ACN. After adding 35 µL 100 mM ABC and Trypsin/LysC 0,1µg/µL stock solution (protein:enzyme ratio of 1:50 (w/w)), samples were incubated and shaken at 37 °C for 17 hours. The reaction was stopped by adding formic acid to a final concentration of 0.1%. Peptide concentrations were determined (Pierce 23290), samples were transferred to a new plate, and frozen at -80 °C until analysis by LC-MS/MS without further conditioning or clean-up.

### Liquid Chromatography–Mass Spectrometry (LC-MS)

450 ng peptides were analyzed on a SCIEX ZenoTOF 7600 System mass spectrometer, coupled to a Waters ACQUITY UPLC M-Class System. Prior to MS analysis, 450 ng sample according to peptide determination was chromatographically separated with a 19 min active gradient, flow 5 µl /min on a Waters HSS T3 column (300 µm x 150 mm, 1.8 µm) heated to 35 °C, where mobile phase A & B are 0.1% formic acid in water and 0.1% formic acid in acetonitrile, respectively. For separation, a gradient from 1-40% B was used. Peptides were detected by a ZenoSWATH MS/MS acquisition scheme with 11 ms accumulation time and 85 variable-size windows. Ion source gas 1 &2 were set as 12 and 60 psi, respectively; Curtain gas 25, CAD gas 7, and source temperature at 150 °C. As source parameters spray voltage was set at 4500 V.

### Protein identification and quantification

Raw data were processed with DIA-NN (version 1.8.1) [26] using the default settings with fragment ion m/z range set from 100-1800, mass accuracy set to 20 ppm and 12 ppm at the MS2 and MS1 level, respectively, scan window set to 7, MBR (Match-between-runs) enabled and Quantification strategy set as “Robust LC (High Precision)”. A spectral library free approach and *mus musculus* UniProt (UP000000589, downloaded on 27.08.2023) were used for annotation. The output was filtered at 1% FDR on the peptide level.

### Metabolic Modeling

Metabolic pathways: the kinetic model comprises the major cellular metabolic pathways of mitochondrial energy metabolism and glycolysis [19]. The model also contains key electrophysiological processes at the inner mitochondrial membrane including the membrane transport of various ions, the mitochondrial membrane potential, and the generation and utilization of the proton-motive force. The time-dependent variations in model variables (i.e., the concentration of metabolites and ions) are governed by first-order differential equations. Time variations of small ions were modeled with kinetic equations of the Goldman–Hodgkin– Katz type. Numerical values for kinetic parameters of the enzymatic rate laws were taken from reported kinetic studies of the isolated enzymes. Maximal enzyme activities (Vmax values) were estimated based on functional characteristics and metabolite concentrations of healthy neuronal tissue [19]. Individual model parametrization: individual metabolic models were established by using the protein intensity profiles delivered via quantitative shotgun proteomics to scale the maximal activities of enzymes and transporters, thereby exploiting the fact that the maximal activity of an enzyme is proportional to the abundance of the enzyme protein according to the following relation:

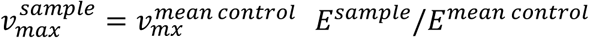

The maximal activities 𝑣^𝑚𝑒𝑎𝑛^ ^𝑐𝑜𝑛𝑡𝑟𝑜𝑙^ for the control were obtained from Berndt et al. [19].

𝐸^𝑚𝑒𝑎𝑛^ ^𝑐𝑜𝑛𝑡𝑟𝑜𝑙^ denotes the mean protein abundance in the control group, and 𝐸^𝑠𝑎𝑚𝑝𝑙𝑒^ denotes the protein abundance of enzyme E in the sample.

Evaluation of energetic capacity: Energetic capacity was assessed under conditions of saturating glucose and oxygen concentrations, which correspond to healthy physiological states. Energetic capacities were evaluated by calculating the changes in the metabolic state induced by an increase in the ATP consumption rate above the resting value. The ATP consumption rate was modeled using a generic hyperbolic rate law

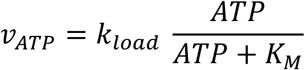

The parameter 𝑘_𝑙𝑜𝑎𝑑_ was stepwise increased until the ATP production rate reached its maximum value.

### Statistical Analysis

Statistical analysis of proteomic data was performed using a two-sample t-test. Significantly regulated proteins between APPPS1 groups and WT controls are indicated as labeled blue dots. PCA plots were made using the Matlab PCA function. For group comparison of metabolic functions, data were checked for normality with a one-sample Kolmogorov Smirnov test. Significant differences between groups were assessed with a two-sided t-test for normally distributed group values; otherwise, the Wilcoxon signed-ranked test was used.

## Results

### Acute isoflurane exposure does not affect glia cell activation

As AD increased the risk of postoperative delirium (POD) [6, 7], 400µm acute brain slices of 11-month-old WT and APPPS1 mice were exposed to 1% followed by 3% isoflurane. Importantly, 1% and 3% isoflurane are clinically relevant concentrations for light and deep anesthesia respectively. The APPPS1 mouse model is a genetic mouse model that represents main characteristics (Aβ deposition and neuroinflammation) of Alzheimer’s disease (AD) [13]. At the age of 11 months, they present extensive Aβ plaques and neuroinflammation [14].

As glia cells react sensitively to changes in the brain, we first assessed the acute effects of isoflurane exposure on microglia and astrocytes. Staining and quantification of microglia and astrocytes confirmed microgliosis and accumulation around the Aβ plaques but no astrogliosis in the APPPS1 mice, while WT mice showed the typical pattern of homeostatic glia cells. No differences between control and isoflurane slices became apparent in both genotypes, indicating that isoflurane treatment did not induce acute cell loss or changes in morphology. In addition, we stained for the AD-related microglial activation marker TREM2 [27]: while WT animals showed almost no TREM2 signal, a co-localization with activated and amoeboid microglia clustered around the Aβ plaques can be observed in APPPS1 mice. However, isoflurane treatment did not result in an altered TREM2 staining intensity.

### Isoflurane decreases CMRO_2_ in brain slices of APPPS1 mice

As AD is associated with impaired energy metabolism [28], we investigated whether neuronal tissue of APPPS1 mice exhibited alterations in energy metabolism under resting conditions and during anesthesia. Brain slices were assessed for oxygen consumption rates under four different conditions: no isoflurane (CTL), 1% isoflurane (1%), 3% isoflurane (3%), and after washout (WO). While no isoflurane refers to spontaneous network activity as described by Bernd et al. [12], 1% isoflurane corresponds to light anesthesia (phase 2) and 3% isoflurane corresponds to deep anesthesia (burst suppression), as assessed by typical electrophysiological patterns, changes in oxidative metabolism, and as described previously [23].

Using Clark electrodes to measure depth profiles of ptiO2 throughout the tissue and applying computational modeling of the reaction-diffusion system to the measured depth profiles, we assessed the cerebral metabolic rates of oxygen (CMRO_2_) in 51 brain slices of 13 WT animals and 43 slices of 11 APPPS1 mice.

Fig. 2A shows the calculated CMRO_2_ of the brain slices under resting conditions without isoflurane (CTL), for light anesthesia (1%), deep anesthesia (3%), and after washout (WO), i.e. 15 minutes after removing the isoflurane from the system through media exchange, for brain slices of WT animals (red, WT) and APPPS1 animals (blue, APPPS1). We found a significant reduction in CMRO_2_ between CTL, 1%, and 3% isoflurane, for both WT and APPPS1 animals, in line with our previous findings in brain slices of Wistar rats [23, 29] that is reversed after washout. We also found a significant reduction in CMRO_2_ between WT and APPPS1 over all conditions (*rmANOVA*: *F*(1, 82) = 5.46, *p* = 0.022), independent of the experimental condition (interaction *p*s > 0.3) and in any given condition (*two-sided t-tests*: CTL: *p* = 0.024, 1%: *p* = 0.043, 3%: *p* = 0.010, WO: *p* = 0.045).

**Fig. 1.**
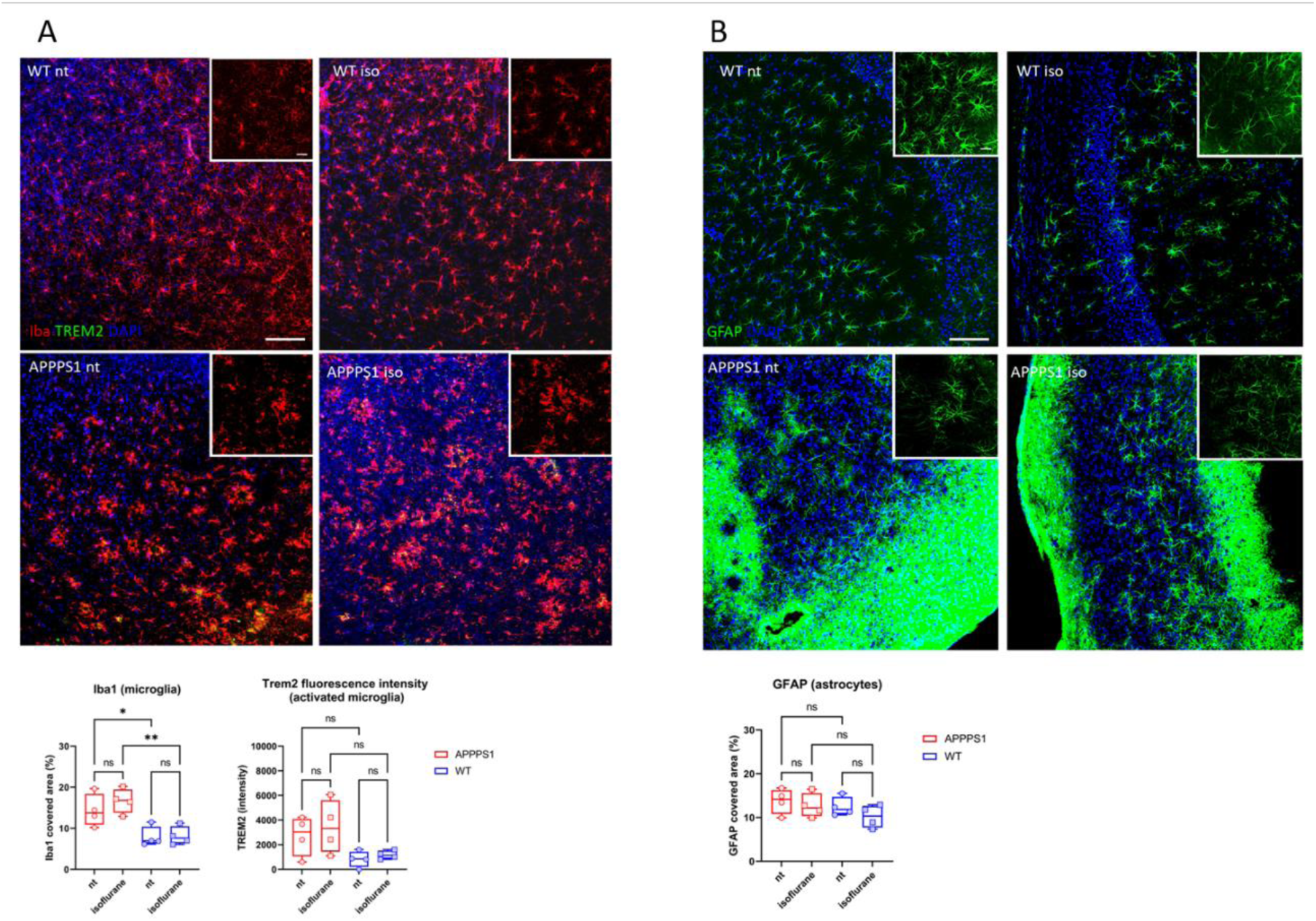
Genotype but not isoflurane affects microglia and astrocytes in WT and APPPS1 brain slices. **(A)** 400 µm brain slices of 11-month-old wild-type (WT) and AD-like APPPS1 mice show microglia (Iba1, red), activated microglia (Trem2, green), and cell nuclei (DAPI, blue). In APPPS1, activated microglia are clustering around amyloid-beta plaques; bar = 100 µm. The insets depict microglial morphology; bar = 25 µm. Quantification of the Iba1 covered area and TREM2 intensity showed microgliosis in APPPS1 mice but no effect of isoflurane. **(B)** 400-µm brain slices of 11-month-old WT and AD-like APPPS1 mice show astrocytes (GFAP, green) and cell nuclei (DAPI, blue); bar = 100 µm. The insets depict astrocytic morphology; bar = 25 µm. Quantification of GFAP covered area showed no astrogliosis in APPPS1 mice and no effect of isoflurane. Group differences were evaluated using one-way ANOVA; significant results are indicated above comparisons; * p <0.05, ** p <0.01).

**Fig. 2.**
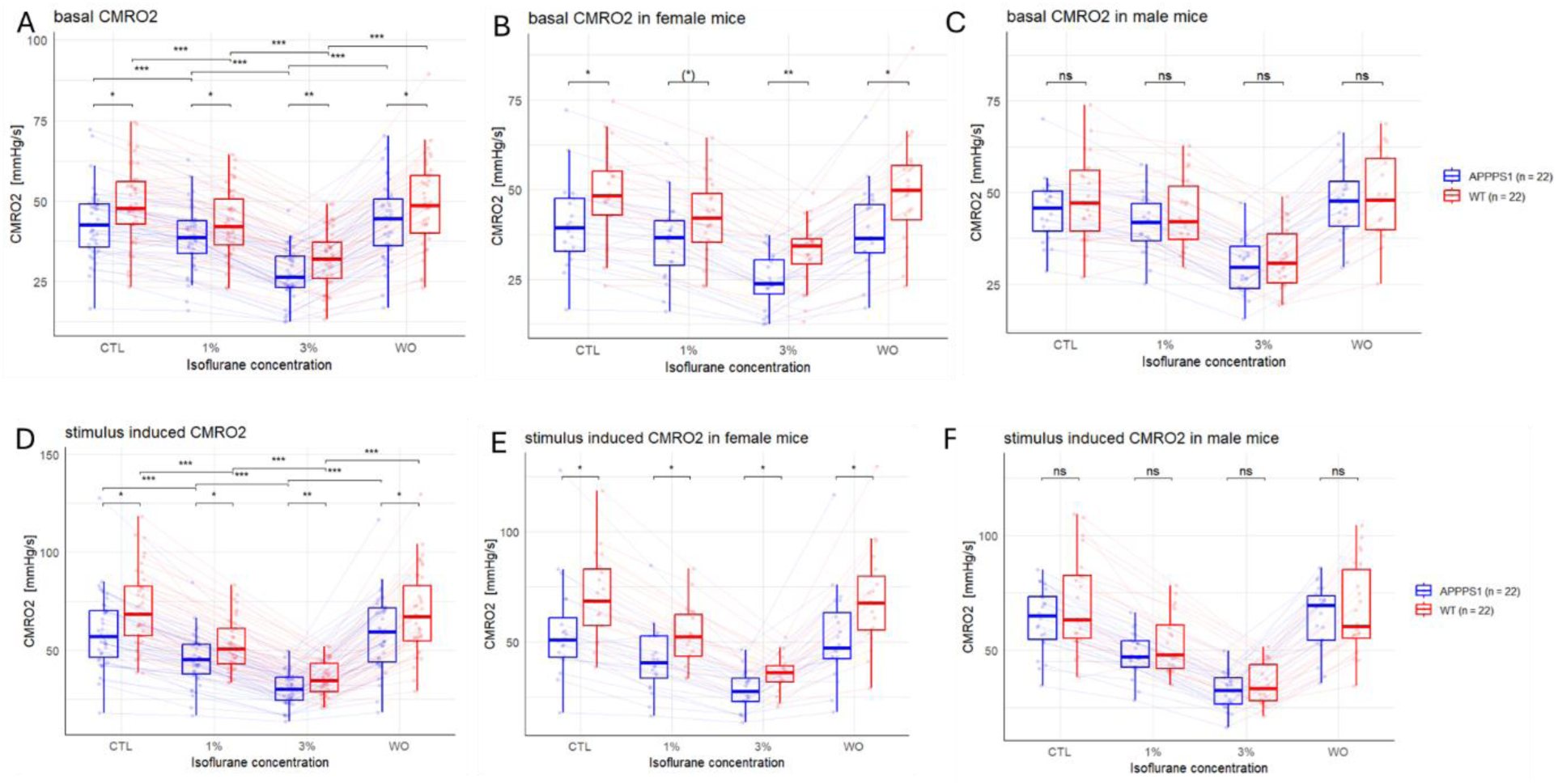
Isoflurane and genotype effects on resting and activity-driven CMRO₂ in WT and APPPS1 brain slices. **(A)** The cerebral metabolic rate of oxygen (CMRO₂) in WT (red) and APPPS1 (blue) brain slices under resting conditions, measured across control (CTL), light anesthesia (1% isoflurane), deep anesthesia (3%), and washout (WO). CMRO₂ significantly decreased with increasing isoflurane concentrations in both genotypes and was reversed upon washout. Across all conditions, APPPS1 slices exhibited significantly lower CMRO₂ than WT slices. **(B+C)** Sex-stratified CMRO₂ under resting conditions. While the isoflurane effect was observed in both sexes, female APPPS1 slices showed significantly lower CMRO₂ than WT females under CTL, 3%, and WO conditions. No significant genotype differences were observed in male slices. **(D)** CMRO₂ during electrical stimulation in the same slices. Stimulation increased CMRO₂ by ∼42% overall under CTL conditions, with a greater increase in WT slices (45%) compared to APPPS1 (39%). Isoflurane significantly suppressed CMRO₂ in both genotypes, consistent with resting data. Across all conditions, APPPS1 slices showed significantly reduced CMRO₂ compared to WT, with a partial attenuation of this difference at 3% isoflurane. **(D+E)** Sex-stratified CMRO₂ during stimulation. Similar to resting conditions, female APPPS1 slices showed significantly lower CMRO₂ than WT females under CTL and WO, with a trend at 1% isoflurane. No significant genotype differences were observed in male slices under any condition. **(A-E)** Group differences were evaluated using Wilcoxon rank-sum or two-sided t-tests, depending on normality; significant results are indicated above comparisons. (*) p <0.01, * p <0.05, ** p <0.01, *** p <0.001).

As sex differences play a role in anesthesia and the development of POD [15] and AD [16], we separated the groups by sex. Fig. 2B+C shows that, while the effect of isoflurane on CMRO_2_ was preserved in both sexes, brain slices from APPPS1 female mice (n = 20) showed a significantly reduced CMRO_2_ compared to WT females (n = 20) across CTL, 3%, and WO, while there was no significant difference in CMRO_2_ for brain slices from male APPPS1 mice (n = 22) compared to brain slices from WT male mice (n = 22), under any condition.

Since energy expenditure is activity-dependent, we also performed experiments with significantly increased energy demand by applying electrical stimulation to the slices as described previously [30]. Figs. 2C and 2D show the corresponding results for the whole experiment and sex-separated groups. Importantly, to minimize heterogeneity due to variability in the slice preparations, all experiments were performed with the same slices.

First, by comparing Figs. 2A and 2D, it is clear that electrical stimulation led to an increase in CMRO_2_ by around 42% overall under CTL conditions, in line with previous results [31], but was slightly higher in WT slices (45%) than in APPPS1 slices (39%), indicating that activity-dependent energy demand was more pronounced in WT animals. Importantly, during stimulation, CMRO2 was significantly higher in WT slices than in APPPS1 slices over all conditions (rmANOVA: F(1, 81) = 7.02, p = 0.010), independent of the experimental condition (interaction ps > 0.08), and for almost every condition separately (two-sided t-tests: CTL: p = 0.011, 1%: p = 0.010, 3%: p = 0.008, WO: p = 0.029). Interestingly, an LMM analysis revealed a significant positive interaction at 3% stimulation (β = 6.00, p = 0.016), suggesting a reduced genotype difference under deep anesthesia conditions. While the simple contrast was not significant (p = 0.146), this may indicate partial compensation in APPPS1 slices at high activation levels. As before, isoflurane significantly reduced CMRO_2_ in both groups (WT: rmANOVA, F(3, 120) = 161.66, p < 2.7e−26, Greenhouse–Geisser corrected; APPPS1: F(3, 123) = 137.66, p < 4.5e−22), with strong pairwise differences between CTL and both 1% and 3% conditions (all p < 0.0001), but not between CTL and WO.

Again, looking at sex-separated groups (Fig. 2E+F), there were clear differences. While there was no significant difference in CMRO_2_ for male animals under any conditions (two-sided t-test: p > 0.05 in all conditions), we found a significantly reduced CMRO_2_ in APPPS1 slices from female animals compared to WT without isoflurane, under 1% and 3% isoflurane and after WO (two-sided t-tests: all p <0.05).

Overall, our results show that there is a decrease in the oxygen consumption rate in brain slices of female but not male APPPS1 mice compared to WT animals and that this difference is more pronounced at higher energy demand, and is attenuated but not leveled by isoflurane administration.

### Isoflurane affects ion homeostasis in APPS1 brain slices

Besides a balanced energy metabolism, maintenance of ion homeostasis is a prerequisite for proper neuronal functionality. AD is associated with altered potassium homeostasis [32, 33].

Therefore, together with p_ti_O_2_, we simultaneously monitored [K^+^]_o_ using ion-sensitive microelectrodes.

Fig. 3A shows [K^+^]_o_ levels under control conditions and upon isoflurane administration and washout for WT and APPPS1 brain slices. Increasing concentrations of isoflurane significantly increased the potassium concentration (AD: rmANOVA, F(3, 90) = 43.42, p < 1.91e-17) as previously shown in Wistar rats [23]. In APPPS1 brain slices, potassium increased from 2.95 mM (CTL) to 3.05 mM (1%, p = 0.7, n.s.) and 3.53 mM (3%, p < 0.001). In WT, levels rose from 3.05 mM (CTL) to 3.28 mM (1%, p = 0.22 n.s.) and 3.83 mM (3%, p < 0.001). There was no significant difference in potassium concentrations in the WT (n = 26) compared to the APPPS1 slices (n = 31) under any condition (rmANOVA > 0.05, two-sided t-tests > 0.05 in all conditions). When separating the groups by sex (Fig. 3B+C), there were significant differences in potassium concentration between APPPS1 slices (male n = 19, female n = 12) and WT slices (male n = 13, female n = 13) for female mice at 3% isoflurane. Additionally, neither genotype nor sex alone had a significant effect on potassium levels. However, there was a significant negative interaction between isoflurane concentration and sex (rmANOVA, *F*(3, 159) = 4.76, p = 0.003). It indicates that the isoflurane effect on potassium concentration differs between males and females with female slices showing a stronger potassium increase at high isoflurane concentrations compared to male slices. Furthermore, a trend towards a three-way interaction among isoflurane concentration, genotype, and sex was observed (rmANOVA, *F*(3, 159) = 2.49, p = 0.06). Specifically, the increase in potassium concentrations was more predominant in WT compared to APPPS1, pointing to a loss of sex-specific differences in potassium homeostasis under isoflurane in the AD-like genotype (Fig. 3B).

**Fig. 3.**
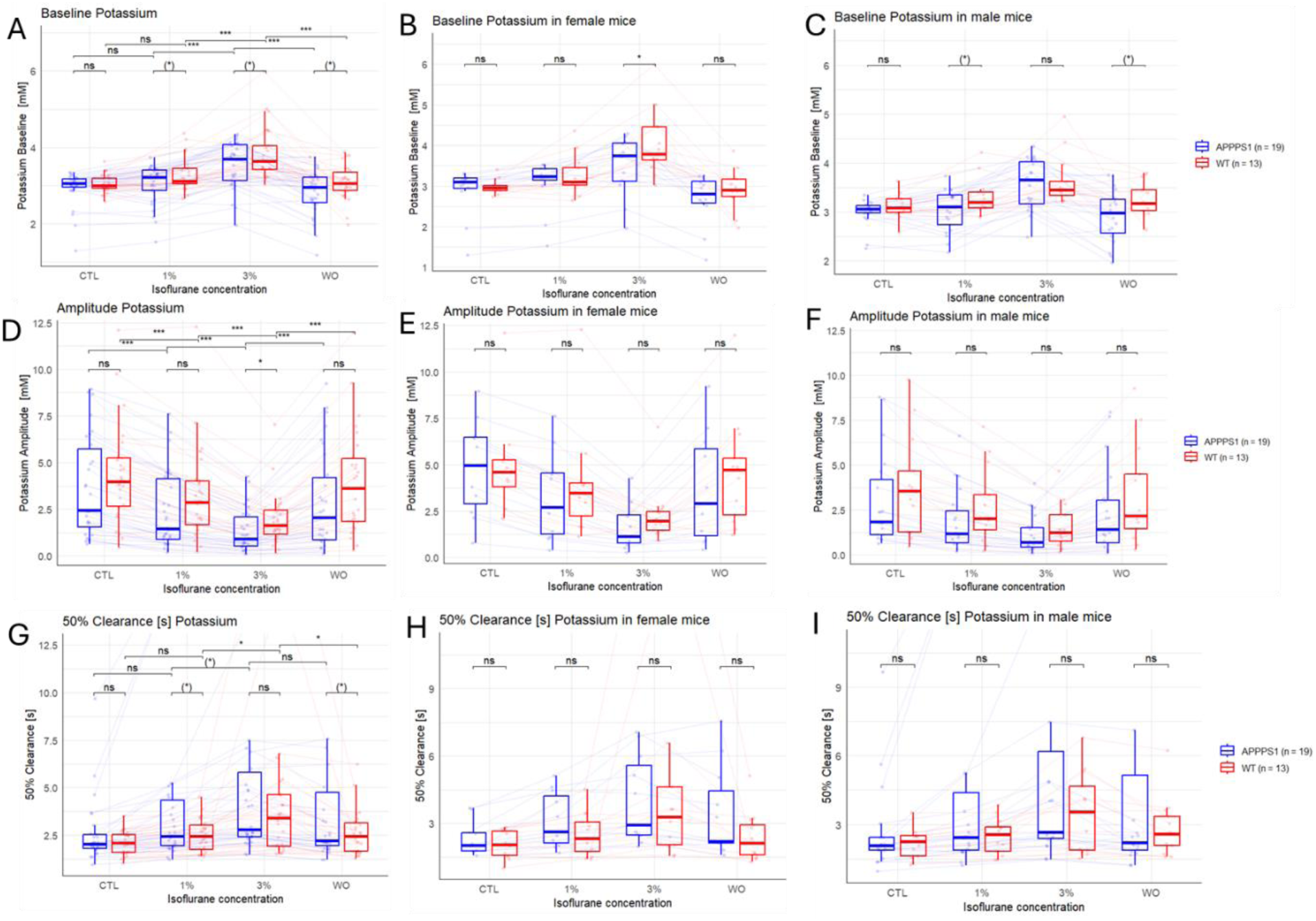
Isoflurane alters extracellular potassium levels and clearance in WT and APPPS1 brain slices. **(A)** Baseline extracellular potassium concentrations [K^+^]_o_ in WT (red) and APPPS1 (blue) brain slices under control conditions (CTL), during isoflurane exposure (1%, 3%), and after washout (WO). Isoflurane significantly increased [K^+^]_o_ in both genotypes in a concentration-dependent and reversible manner. **(B+C)** Sex-specific effects of isoflurane on [K^+^]_o_ at rest in female (B) and male (C) slices. While no main effects of sex or genotype were detected, a significant interaction between sex and isoflurane concentration was observed, with female slices showing a stronger increase in [K^+^]_o_ at higher isoflurane concentrations, particularly in WT animals. **(D)** Peak [K^+^]_o_ during stimulation under increasing isoflurane concentrations. Isoflurane reduced stimulus-induced potassium elevations in both genotypes, with a significantly greater reduction in APPPS1 slices at 3% isoflurane. **(E+F)** Sex-stratified peak [K^+^]_o_ responses to stimulation in female (E) and male (F) slices. A trend for sex-dependent differences was observed: males showed greater reduction at 1% and females at 3% isoflurane, suggesting a potential sex-dependent sensitivity to isoflurane. **(G)** Isoflurane effects on potassium clearance dynamics. Time to 50% recovery (T50) of stimulus-induced [K^+^]_o_ was prolonged with increasing isoflurane concentrations in both genotypes, indicating slower potassium clearance. **(H+I)** T50 clearance time stratified by sex in female (H) and male (I) slices. APPPS1 females exhibited significantly prolonged T50 at 3% isoflurane. Differences in potassium clearance were more pronounced in males than females at lower isoflurane concentrations but diminished at 90% clearance levels (not shown). **(A-I)** Group differences were evaluated using Wilcoxon rank-sum or two-sided t-tests, depending on normality; significant results are indicated above comparisons. (*) p <0.01, * p <0.05, ** p <0.01, *** p <0.001).

We also measured the effects of stimulus-induced increase in extracellular potassium during isoflurane administration. Increasing concentrations of isoflurane lead to a decrease in potassium amplitude in response to stimulation in WT and APPPS1 slices (Fig. 3D). The stimulus-induced [K^+^]_o_ increase significantly declined with isoflurane application (AD: rmANOVA, F(3, 87) = 38.98, p < 4.61e-16) (Fig. 3D). In APPPS1, the stimulus-induced [K^+^]_o_ increase declined from 3.73 mM (CTL) to 2.42 mM (1%, p < 0.001) and 1.33 mM (3%, p < 0.001).

In WT, amplitudes declined from 4.32 mM (CTL) to 3.27 mM (1%, p < 0.001) and 1.12 mM (3%, p < 0.001). We also found a significant decrease in stimulus-induced potassium concentrations in APPPS1 compared to WT animals under 3% isoflurane (Wilcoxon rank test, p = 0.034). Sex-differentiated groups showed no significant differences under any condition (Fig. 3 E+F).

Although neither genotype (rmANOVA*; F*(1, 52) = 1.91, *p* = 0.173) nor sex alone (rmANOVA*; F*(1, 52) = 3.74, *p* = 0.059) showed significant main effects, there was a tendency for sex differences. Furthermore, a trend toward an interaction between isoflurane concentration and sex was observed (*F*(3, 156) = 2.59, *p* = 0.055), suggesting possible sex-related modulation of the isoflurane effect on amplitude. Specifically, male animals tended to show a stronger reduction in amplitude at lower isoflurane concentrations (1%), while female animals exhibited a more pronounced reduction at higher concentrations (3%), indicating potential sex-dependent differences in isoflurane sensitivity.

To distinguish whether the observed differences in potassium concentrations are a result of altered potassium release or indicative of a potential difference in ion pumping after stimulation, we monitored the clearance of potassium, e.g. the regression of potassium concentration back to baseline levels after stimulations. Proportionally to an increase in isoflurane concentration, the T50 (AD: rmANOVA, F(3, 87) = 9.25, p < 0.001; WT: F(3, 75) = 6.42, p < 0.001) (Fig. 3G) and T90 (AD: rmANOVA, F(3, 78) = 29.3, p < 0.001; WT: F(3, 72) = 34.65, p < 0.001) (not shown) prolonged, respectively. The relative percentage (deltaK) of the stimulus-induced rise in [K^+^]_o_ remaining after 1, 2, and 5 s increased with increasing isoflurane percentage (rmANOVA p < 0.05 in all, independent of genotype) (not shown). While there was a tendency for slower potassium clearance at the 50% level for APPPS1 compared to WT animals at 1% isoflurane (two-sided t-test, p = 0.058) and WO (p = 0.051), these differences disappeared at 90% and higher clearance levels. These differences in potassium clearance seemed to be more pronounced in male than in female mice (Fig. 3H+I).

### Proteomic characterization

To discover the molecular and proteomic alterations underlying the observed physiological changes, we investigated the proteome of the brain slices by mass spectrometry in WT and APPPS1 mice. Overall, we identified 5,897 proteins. The volcano plot in Fig. 4A illustrates differences between the mean protein intensities of APPPS1 and WT mice. We found 36 proteins significantly upregulated in APPPS1 mice, but no downregulated proteins (p-value <0.05 and log2-fold change >2).

**Fig. 4.**
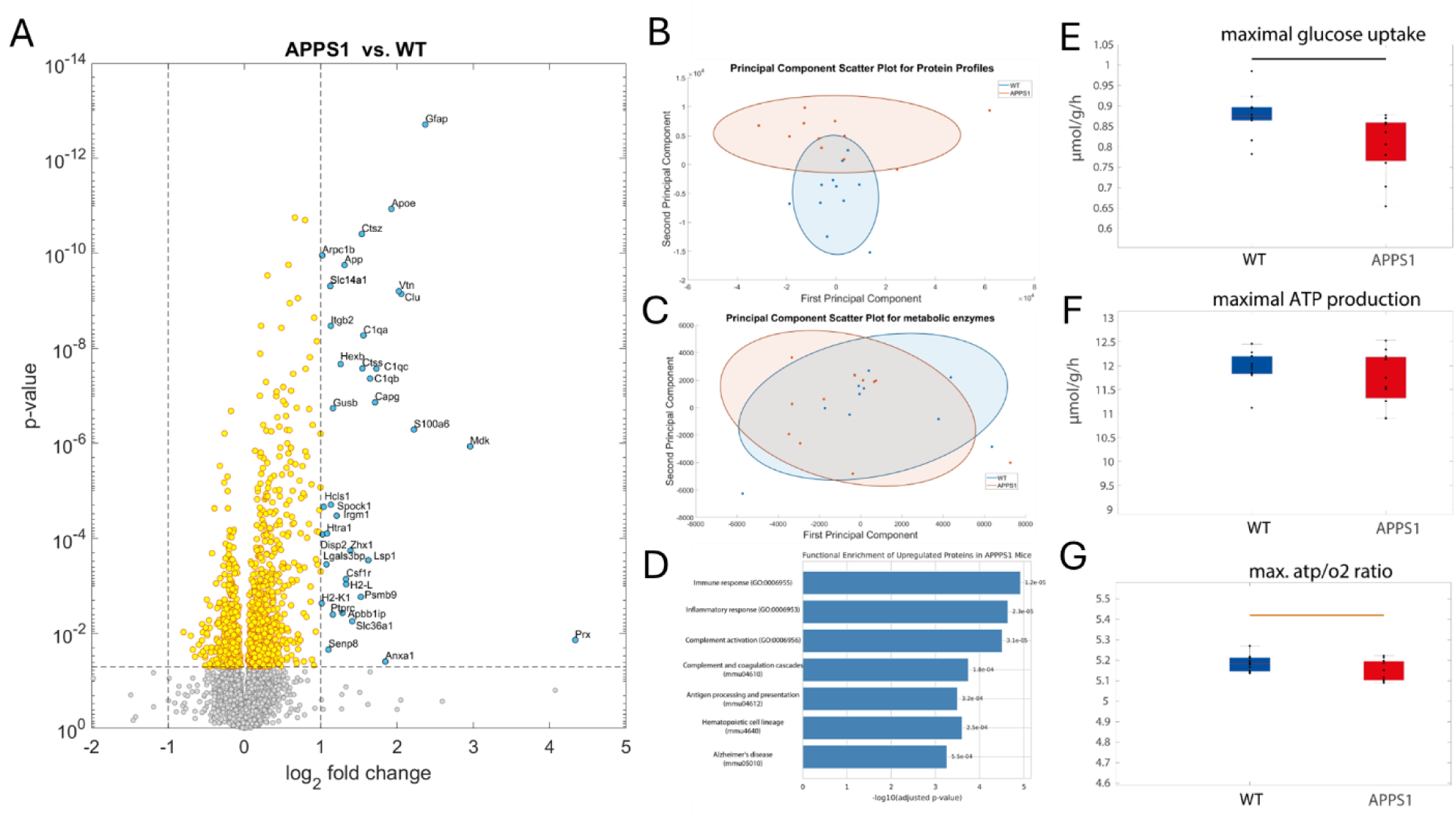
Proteomic and metabolic changes in APPPS1 brain slices reflect Alzheimer’s-related neuroinflammation and preserved energy production capacity. (A) Volcano plot displaying the log₂ fold changes of 5,897 proteins quantified in brain slices from APPPS1 mice (n = 11) versus WT controls (n = 11). Proteins on the left are downregulated, while those on the right are upregulated in APPPS1 mice. A total of 37 proteins were significantly upregulated (p < 0.05; log₂ fold change > 1), shown in blue. (B) Principal component analysis (PCA) of all detected proteins shows a clear separation between WT and APPPS1 groups, indicating broad proteomic alterations. (C) PCA restricted to metabolic enzymes does not reveal a clear group separation, suggesting preserved overall metabolic protein expression. (D) Pathway enrichment analysis of the 36 upregulated proteins reveals significant enrichment of biological processes related to immune activation—including “immune response,” “inflammatory response,” and “complement activation”—and KEGG pathways such as “complement and coagulation cascades,” “antigen processing and presentation,” and “Alzheimer’s disease,” indicating upregulation of disease-relevant molecular functions. (E-G) Functional metabolic modeling based on the proteomic data was used to simulate neuronal energy metabolism. While maximal glucose uptake (E) and ATP production capacity (F) were maintained in APPPS1 slices, glycolytic flux was significantly reduced (p = 0.02), and the ATP/O₂ ratio (G) showed a trend toward decreased energetic efficiency (p = 0.058). Statistical comparisons were made using Wilcoxon rank-sum or two- sided t-tests, as appropriate; significance levels are indicated by brackets above group comparisons (black: p <0.05; orange: p <0.01).

Functional grouping indicated that several of these proteins are directly involved in amyloid metabolism (APP, APOE, CLU), while others mark reactive astrocytes (GFAP, S100A6), and disease-associated microglia (C1qa/b/c, CSF1R, HEXB, CTSS). Additional proteins were implicated in antigen presentation (H2-K1, PSMB9, IRGM1), oxidative stress (PRX), and extracellular matrix remodeling (SPOCK1, HTRA1). These findings reflect key features of Alzheimer’s pathology, including innate immune activation, glial remodeling, and altered proteostasis, and support the utility of the APPPS1 model for studying neuroinflammatory and amyloidogenic processes.

To investigate the functional landscape of proteins associated with neurodegeneration in the APPPS1 AD-like mouse model, we performed a comprehensive enrichment analysis of these 37 proteins using the g:Profiler platform. The APPPS1 model overexpresses a mutant form of the human amyloid precursor protein (APP) and is characterized by extracellular amyloid plaque deposition, neuroinflammation, and reactive gliosis.

Gene Ontology (GO) enrichment analysis revealed a significant overrepresentation of biological processes related to immune activation. Highly enriched GO terms included “immune response” (GO:0006955, adj. p = 1.2e-5), “inflammatory response” (GO:0006954, adj. p = 2.3e-5), and “complement activation” (GO:0006956, adj. p = 3.1e-5), consistent with known microglial and complement-mediated synaptic pruning in AD pathology.

KEGG pathway analysis identified “complement and coagulation cascades” (mmu04610, adj. p = 1.8e-4), “hematopoietic cell lineage” (mmu04640, adj. p = 2.5e-4), and “antigen processing and presentation” (mmu04612, adj. p = 3.2e-4) as significantly enriched. Notably, the “Alzheimer’s disease” pathway (mmu05010, adj. p = 5.5e-4) was also represented, underscoring the disease relevance of the selected proteins (see Fig. 4D).

Notably, none of these proteins was associated with changes in energy metabolism. However, the observed alterations in oxygen consumption hint at possible metabolic alterations that potentially also alter the ability to generate the energy needed for proper neuronal functionality. To specifically analyze metabolic alterations, we selected the subset of metabolic proteins and performed separate PCA analysis. While the PCA plot considering all proteins showed a separation of the WT and APPPS1 groups (Fig. 4B), the metabolic proteins did not show any separation (Fig. 4C). Next, we integrated protein abundance data of metabolic enzymes into a molecular resolved kinetic model of neuronal metabolism [19] and simulated the maximal ATP as described by Liotta et al. [34]. As shown in Fig. 4 (right panels), we found no difference in the maximal ATP production capacities in the APPPS1 group. However, we found a significant reduction of glycolysis (two-sided t-test, p= 0.02) in the APPPS1 mice and a tendency toward reduced efficiency as assessed by ATP/O_2_ ratio (two- sided t-test, p = 0.058).

## Discussion

Anesthesia is a major risk factor for postoperative complications like POD. However, the underlying molecular mechanisms are still unclear. This study aimed to identify neurophysiological and metabolic alterations in the brain of the APPPS1 AD-like mouse model and to assess their interaction with anesthetic exposure. By combining oxygen consumption profiling, potassium ion dynamics, glial immunostaining, and proteomic analyses, we delineated a multifaceted picture of AD-related dysfunction, particularly highlighting sex-specific vulnerabilities.

### Energy Metabolism Is Impaired in Female APPPS1 Mice

Our data demonstrate that energy metabolism is compromised in APPPS1 brain tissue, with this impairment most evident under increased energetic demand. While baseline CMRO₂ (oxygen consumption) was reduced in AD-like animals compared to WT under all conditions, the reduction became more prominent during electrical stimulation. This result correlates with oxygen measurements of human iPSC-derived neurons with APP and PS1 mutations [35], which also show less oxygen consumption rate under basal conditions. Furthermore, a decreased oxygen consumption rate in immortalized astrocytes from the AD-like mouse model 3Tg [36] and in mitochondria isolated from the hearts of the AD-like mouse model APPSWE/PS1 Tg [37] was observed under basal conditions. The underlying reason is most probably the often-observed mitochondrial dysfunction in AD-like mouse models and also human AD patients [9]. This suggests that while metabolic compensation may mask deficits at rest, energetic challenges unmask bioenergetic limitations in the AD model.

Sex emerged as a strong modulator of this effect: female APPPS1 mice exhibited significantly reduced CMRO_2_ compared to WT across both resting and stimulated states. In contrast, no significant CMRO_2_ difference was observed between male APPPS1 and WT mice. This aligns with clinical findings that women are at increased risk for AD progression [16]. Interestingly, POD is more prevalent in men [17, 18], although the reasons are not clear yet. Higher prevalence of comorbidities in men (e.g., cardiovascular disease, substance use) and/or potential underdiagnosis in females due to subtler symptom presentation could contribute to this disparity. Different sex-specific mechanisms—potentially including estrogen-driven mitochondrial regulation, differential expression of metabolic enzymes, and immune responsiveness—may underlie these findings and warrant targeted investigation.

### Anesthetic Exposure Equalizes Metabolic Output

As expected, isoflurane exposure significantly suppressed CMRO₂ across all genotypes and sexes consistent in line with our previous research [23, 29, 31]. Importantly, it maintains metabolic differences between groups, even under deep anesthesia (3%). This suggests a “metabolic floor” effect where neural activity and associated oxygen demands are uniformly reduced. While this suppressive effect was reversible after washout, indicating no long-term metabolic impairment under acute exposure in this *ex vivo* model, metabolic differences between WT and APPPS1 slices were preserved. Our data suggest an impaired oxidative capacity of the APPPS1 mice which was preserved under anesthesia. This could result in hypoxia, either globally or in distinct brain regions, which could drive AD-relevant Aβ and tau deposition, similar to patients with obstructive sleep apnea (OSA) [38]. Furthermore, hypoxia impairs ATP production, potentially compromising synaptic transmission and cellular activity of the different CNS-residential cells. For example, phagocytosis—the removal of dead cells, synapses, myelin debris, and Aβ—is an important function of microglia phagocytosis for general “brain hygiene”. Whether this holds *in vivo*—especially under prolonged or repeated anesthetic exposure—remains a critical question.

### Potassium homeostasis Shows Subtle Phenotype Differences

[K⁺]ₒ dynamics, as a surrogate for neuronal excitability and energy-dependent ion regulation, also reflected differences between genotypes and sexes. Isoflurane induced expected increases in resting [K⁺]ₒ and reductions in evoked field potentials, consistent with reduced neuronal responsiveness [29].

While no clear genotype-dependent difference was observed in baseline or stimulated [K⁺]ₒ levels, female/male APPPS1 mice showed more pronounced divergence from WT in specific conditions, supporting a pattern of sex-specific vulnerability. Potassium clearance kinetics (T50) were slowed by isoflurane and also tended to be slower in APPS1 mice compared to WT mice, implying that part of the deficit may lie in the energetic capacity to support stimulus-induced K⁺ clearance mechanisms.

### Glial Reactivity: A Primed, Not Provoked, Landscape

Our immunohistological analysis confirmed microglial activation in APPPS1 mice, particularly in proximity to Aβ plaques, along with increased expression of the activation marker TREM2. However, no astrogliosis was observed, and isoflurane exposure did not exacerbate glial activation. These findings suggest that in this acute *ex vivo* context, glia in the APPPS1 brain are already in a “primed” state due to chronic pathology but do not further react to acute anesthetic exposure.

In contrast to other *in vivo* models showing glial activation after repeated or chronic anesthesia [39, 40], our results emphasize that both exposure duration and biological context influence neuroimmune responses. Given the strong association between microglial activation, synaptic pruning, phagocytosis, and cognitive decline in AD, the preexisting glial state in APPPS1 mice may sensitize neural circuits to additional stressors, even if the glial cells themselves do not show acute hyperreactivity to anesthesia.

### Proteomic Analysis Supports Immune Activation Reveals Glycolytic Weakness

Proteomic profiling revealed immune and inflammatory signatures in APPPS1 tissue, consistent with known AD hallmarks. Upregulated proteins involved in complement activation, Aβ processing, and glial reactivity reaffirm the model’s disease relevance. However, no metabolic enzymes were differentially expressed at the protein level. PCA based on metabolic proteins failed to separate genotypes, supporting the idea that metabolic dysfunction arises from regulatory, post-translational, or subcellular mechanisms rather than gross protein abundance changes.

Integration of protein abundance data into a molecularly detailed kinetic model of neuronal metabolism revealed functional impairments in APPPS1 mice. Notably, a significant reduction in glycolytic activity under conditions of maximal ATP demand was observed (two-sided t-test, *p* = 0.02), accompanied by a trend toward reduced metabolic efficiency, as indicated by a lower ATP/O₂ yield (*p* = 0.058). These results suggest that even if total ATP production remains relatively stable, the architecture of the metabolic network may still be compromised.

Although approximately 90% of neuronal ATP is typically generated via oxidative phosphorylation [41], glycolysis plays a critical role in rapid ATP buffering and the localized fueling of energy-intensive processes. A reduction in glycolytic flux could therefore undermine the cell’s ability to meet sudden or spatially constrained energy demands, particularly under high activity conditions. Essential ATP-dependent functions—such as ion homeostasis and synaptic signaling—may be especially vulnerable in this context.

Moreover, glycolytic ATP is known to preferentially support specific cellular processes, such as calcium handling [42]. Astrocytes, in particular, depend heavily on glycolysis to maintain ion gradients and clear neurotransmitters [43]. As such, even modest impairments in glycolytic capacity could selectively compromise these functions. This type of selective vulnerability may lead to disrupted intercellular communication, impaired homeostasis, and ultimately, degradation of coordinated neural network activity in the APPPS1 brain.

### Conclusion and Outlook

In summary, our study shows that AD-like APPPS1 mice, particularly females, exhibit subtle yet functionally important deficits in neuronal energy metabolism—especially under conditions of increased demand. Isoflurane anesthesia reduces metabolic differences acutely but may conceal underlying vulnerabilities and could induce hypoxia. Potassium homeostasis is subtly impaired, potentially reflecting energetic limitations tied to glycolytic dysfunction. Glial cells are chronically activated in the AD model but do not exhibit acute responsiveness to anesthesia in the slice preparation.

These findings provide mechanistic insights into the heightened perioperative vulnerability of AD patients and underscore the importance of considering sex as a biological variable. Future *in vivo* studies should investigate whether chronic anesthetic exposure exacerbates these vulnerabilities and test whether metabolic support strategies—such as glycolytic enhancement—can mitigate POD risk in AD models.

## Notes

### Competing Interest Statement

The authors have declared no competing interest.

### Summary of Updates

This version of the manuscript has been revised to update the following: The Materials and Methods section has been revised to correct and replace the previously incorrect text concerning proteomics. The updated version now includes accurate and detailed descriptions under the subsections Proteomics Sample Preparation, Liquid Chromatography-Mass Spectrometry (LC-MS), and Protein Identification and Quantification. These revisions were necessary because the initial submission contained placeholder or incorrect content that did not accurately reflect the experimental procedures used in the study. The new text provides a clear and complete account of the methods employed for tissue preparation, peptide processing, mass spectrometry acquisition, and subsequent protein identification and quantification. This correction ensures the methodological accuracy and reproducibility of the proteomic data presented in the manuscript.

## References

1. Inouye SK, Marcantonio ER, Kosar CM, Tommet D, Schmitt EM, Travison TG, Saczynski JS, Ngo LH, Alsop DC, Jones RN, The short-term and long-term relationship between delirium and cognitive trajectory in older surgical patients. Alzheimers Dement, (2016). 12(7): p. 766–75. DOI: 10.1016/j.jalz.2016.03.005.

2. Maldonado JR, Delirium pathophysiology: An updated hypothesis of the etiology of acute brain failure. Int J Geriatr Psychiatry, (2018). 33(11): p. 1428–1457. DOI: 10.1002/gps.4823.

3. Dunn J Grider MH, Physiology, Adenosine Triphosphate, in StatPearls. 2025: Treasure Island (FL).

4. Shigetomi E, Sakai K, Koizumi S, Extracellular ATP/adenosine dynamics in the brain and its role in health and disease. Front Cell Dev Biol, (2023). 11: p. 1343653. DOI: 10.3389/fcell.2023.1343653.

5. Di Virgilio F, Vultaggio-Poma V, Falzoni S, Giuliani AL, Extracellular ATP: A powerful inflammatory mediator in the central nervous system. Neuropharmacology, (2023). 224: p. 109333. DOI: 10.1016/j.neuropharm.2022.109333.

6. An Z, Xiao L, Chen C, Wu L, Wei H, Zhang X, Dong L, Analysis of risk factors for postoperative delirium in middle-aged and elderly fracture patients in the perioperative period. Sci Rep, (2023). 13(1): p. 13019. DOI: 10.1038/s41598-023-40090-z.

7. Needham MJ, Webb CE, Bryden DC, Postoperative cognitive dysfunction and dementia: what we need to know and do. Br J Anaesth, (2017). 119(suppl_1): p. i115– i125. DOI: 10.1093/bja/aex354.

8. D’Alessandro MCB, Kanaan S, Geller M, Pratico D, Daher JPL, Mitochondrial dysfunction in Alzheimer’s disease. Ageing Res Rev, (2025). 107: p. 102713. DOI: 10.1016/j.arr.2025.102713.

9. McGill Percy KC, Liu Z, Qi X, Mitochondrial dysfunction in Alzheimer’s disease: Guiding the path to targeted therapies. Neurotherapeutics, (2025). 22(3): p. e00525. DOI: 10.1016/j.neurot.2025.e00525.

10. Cao S, Wu Y, Gao Z, Tang J, Xiong L, Hu J, Li C, Automated phenotyping of postoperative delirium-like behaviour in mice reveals the therapeutic efficacy of dexmedetomidine. Commun Biol, (2023). 6(1): p. 807. DOI: 10.1038/s42003-023-05149-7.

11. Davis DHJ, Skelly DT, Murray C, Hennessy E, Bowen J, Norton S, Brayne C, Rahkonen T, Sulkava R, Sanderson DJ, Rawlins JN, Bannerman DM, MacLullich AMJ, Cunningham C, Worsening cognitive impairment and neurodegenerative pathology progressively increase risk for delirium. Am J Geriatr Psychiatry, (2015). 23(4): p. 403– 415. DOI: 10.1016/j.jagp.2014.08.005.

12. Berndt N, Rosner J, Haq RU, Kann O, Kovacs R, Holzhutter HG, Spies C, Liotta A, Possible neurotoxicity of the anesthetic propofol: evidence for the inhibition of complex II of the respiratory chain in area CA3 of rat hippocampal slices. Arch Toxicol, (2018). 92(10): p. 3191–3205. DOI: 10.1007/s00204-018-2295-8.

13. Radde R, Bolmont T, Kaeser SA, Coomaraswamy J, Lindau D, Stoltze L, Calhoun ME, Jaggi F, Wolburg H, Gengler S, Haass C, Ghetti B, Czech C, Holscher C, Mathews PM, Jucker M, Abeta42-driven cerebral amyloidosis in transgenic mice reveals early and robust pathology. EMBO Rep, (2006). 7(9): p. 940–6. DOI: 10.1038/sj.embor.7400784.

14. Freitag K, Sterczyk N, Wendlinger S, Obermayer B, Schulz J, Farztdinov V, Mulleder M, Ralser M, Houtman J, Fleck L, Braeuning C, Sansevrino R, Hoffmann C, Milovanovic D, Sigrist SJ, Conrad T, Beule D, Heppner FL, Jendrach M, Spermidine reduces neuroinflammation and soluble amyloid beta in an Alzheimer’s disease mouse model. J Neuroinflammation, (2022). 19(1): p. 172. DOI: 10.1186/s12974-022-02534-7.

15. DiMaria S, Mangano N, Bruzzese A, Bartula B, Parikh S, Costa A, Genetic Variation and Sex-Based Differences: Current Considerations for Anesthetic Management. Curr Issues Mol Biol, (2025). 47(3). DOI: 10.3390/cimb47030202.

16. Lopez-Lee C, Torres ERS, Carling G, Gan L, Mechanisms of sex differences in Alzheimer’s disease. Neuron, (2024). 112(8): p. 1208–1221. DOI: 10.1016/j.neuron.2024.01.024.

17. Wang H, Guo X, Zhu X, Li Y, Jia Y, Zhang Z, Yuan S, Yan F, Gender Differences and Postoperative Delirium in Adult Patients Undergoing Cardiac Valve Surgery. Front Cardiovasc Med, (2021). 8: p. 751421. DOI: 10.3389/fcvm.2021.751421.

18. Wittmann M, Kirfel A, Jossen D, Mayr A, Menzenbach J, The Impact of Perioperative and Predisposing Risk Factors on the Development of Postoperative Delirium and a Possible Gender Difference. Geriatrics (Basel), (2022). 7(3). DOI: 10.3390/geriatrics7030065.

19. Berndt N, Kann O, Holzhutter HG, Physiology-based kinetic modeling of neuronal energy metabolism unravels the molecular basis of NAD(P)H fluorescence transients. J Cereb Blood Flow Metab, (2015). 35(9): p. 1494–506. DOI: 10.1038/jcbfm.2015.70.

20. Angamo EA, Rosner J, Liotta A, Kovacs R, Heinemann U, A neuronal lactate uptake inhibitor slows recovery of extracellular ion concentration changes in the hippocampal CA3 region by affecting energy metabolism. J Neurophysiol, (2016). 116(5): p. 2420–2430. DOI: 10.1152/jn.00327.2016.

21. Huchzermeyer C, Berndt N, Holzhutter HG, Kann O, Oxygen consumption rates during three different neuronal activity states in the hippocampal CA3 network. J Cereb Blood Flow Metab, (2013). 33(2): p. 263–71. DOI: 10.1038/jcbfm.2012.165.

22. Honemann CW, Washington J, Honemann MC, Nietgen GW, Durieux ME, Partition coefficients of volatile anesthetics in aqueous electrolyte solutions at various temperatures. Anesthesiology, (1998). 89(4): p. 1032–5. DOI: 10.1097/00000542-199810000-00032.

23. Reiffurth C, Berndt N, Gonzalez Lopez A, Schoknecht K, Kovacs R, Maechler M, Grote Lambers M, Dreier JP, Friedman A, Spies C, Liotta A, Deep Isoflurane Anesthesia Is Associated with Alterations in Ion Homeostasis and Specific Na+/K+-ATPase Impairment in the Rat Brain. Anesthesiology, (2023). 138(6): p. 611–623. DOI: 10.1097/ALN.0000000000004553.

24. Kasischke KA, Lambert EM, Panepento B, Sun A, Gelbard HA, Burgess RW, Foster TH, Nedergaard M, Two-photon NADH imaging exposes boundaries of oxygen diffusion in cortical vascular supply regions. J Cereb Blood Flow Metab, (2011). 31(1): p. 68–81. DOI: 10.1038/jcbfm.2010.158.

25. Muller T, Kalxdorf M, Longuespee R, Kazdal DN, Stenzinger A, Krijgsveld J, Automated sample preparation with SP3 for low-input clinical proteomics. Mol Syst Biol, (2020). 16(1): p. e9111. DOI: 10.15252/msb.20199111.

26. Demichev V, Messner CB, Vernardis SI, Lilley KS, Ralser M, DIA-NN: neural networks and interference correction enable deep proteome coverage in high throughput. Nat Methods, (2020). 17(1): p. 41–44. DOI: 10.1038/s41592-019-0638-x.

27. Keren-Shaul H, Spinrad A, Weiner A, Matcovitch-Natan O, Dvir-Szternfeld R, Ulland TK, David E, Baruch K, Lara-Astaiso D, Toth B, Itzkovitz S, Colonna M, Schwartz M, Amit I, A Unique Microglia Type Associated with Restricting Development of Alzheimer’s Disease. Cell, (2017). 169(7): p. 1276–1290 e17. DOI: 10.1016/j.cell.2017.05.018.

28. Yu L, Jin J, Xu Y, Zhu X, Aberrant Energy Metabolism in Alzheimer’s Disease. J Transl Int Med, (2022). 10(3): p. 197–206. DOI: 10.2478/jtim-2022-0024.

29. Schoknecht K, Maechler M, Wallach I, Dreier JP, Liotta A, Berndt N, Isoflurane lowers the cerebral metabolic rate of oxygen and prevents hypoxia during cortical spreading depolarization in vitro: An integrative experimental and modeling study. J Cereb Blood Flow Metab, (2024). 44(6): p. 1000–1012. DOI: 10.1177/0271678X231222306.

30. Liotta A, Rosner J, Huchzermeyer C, Wojtowicz A, Kann O, Schmitz D, Heinemann U, Kovacs R, Energy demand of synaptic transmission at the hippocampal Schaffer-collateral synapse. J Cereb Blood Flow Metab, (2012). 32(11): p. 2076–83. DOI: 10.1038/jcbfm.2012.116.

31. Berndt N, Kovacs R, Schoknecht K, Rosner J, Reiffurth C, Maechler M, Holzhutter HG, Dreier JP, Spies C, Liotta A, Low neuronal metabolism during isoflurane-induced burst suppression is related to synaptic inhibition while neurovascular coupling and mitochondrial function remain intact. J Cereb Blood Flow Metab, (2021). 41(10): p. 2640–2655. DOI: 10.1177/0271678X211010353.

32. Roberts BR, Doecke JD, Rembach A, Yevenes LF, Fowler CJ, McLean CA, Lind M, Volitakis I, Masters CL, Bush AI, Hare DJ, group Ar, *Rubidium and potassium levels are altered in Alzheimer’s disease brain and blood but not in cerebrospinal fluid*. Acta Neuropathol Commun, (2016). 4(1): p. 119. DOI: 10.1186/s40478-016-0390-8.

33. Mahan B, Hu Y, Lahoud E, Nestmeyer M, McCoy-West A, Manestar G, Fowler C, Bush AI, Moynier F, Stable potassium isotope ratios in human blood serum towards biomarker development in Alzheimer’s disease. Metallomics, (2024). 16(9). DOI: 10.1093/mtomcs/mfae038.

34. Liotta A, Loroch S, Wallach I, Klewe K, Marcus K, Berndt N, Metabolic Adaptation in Epilepsy: From Acute Response to Chronic Impairment. Int J Mol Sci, (2024). 25(17). DOI: 10.3390/ijms25179640.

35. MacMullen C, Sharma N, Davis RL, Mitochondrial dynamics and bioenergetics in Alzheimer’s induced pluripotent stem cell-derived neurons. Brain, (2025). 148(4): p. 1405–1420. DOI: 10.1093/brain/awae364.

36. Dematteis G, Vydmantaite G, Ruffinatti FA, Chahin M, Farruggio S, Barberis E, Ferrari E, Marengo E, Distasi C, Morkuniene R, Genazzani AA, Grilli M, Grossini E, Corazzari M, Manfredi M, Lim D, Jekabsone A, Tapella L, Proteomic analysis links alterations of bioenergetics, mitochondria-ER interactions and proteostasis in hippocampal astrocytes from 3xTg-AD mice. Cell Death Dis, (2020). 11(8): p. 645. DOI: 10.1038/s41419-020-02911-1.

37. Aishwarya R, Abdullah CS, Remex NS, Bhuiyan MAN, Lu XH, Dhanesha N, Stokes KY, Orr AW, Kevil CG, Bhuiyan MS, Diastolic dysfunction in Alzheimer’s disease model mice is associated with Abeta-amyloid aggregate formation and mitochondrial dysfunction. Sci Rep, (2024). 14(1): p. 16715. DOI: 10.1038/s41598-024-67638-x.

38. Bhuniya S, Goyal M, Chowdhury N, Mishra P, Intermittent hypoxia and sleep disruption in obstructive sleep apnea increase serum tau and amyloid-beta levels. J Sleep Res, (2022). 31(5): p. e13566. DOI: 10.1111/jsr.13566.

39. Kho W, von Haefen C, Paeschke N, Nasser F, Endesfelder S, Sifringer M, Gonzalez-Lopez A, Lanzke N, Spies CD, Dexmedetomidine Restores Autophagic Flux, Modulates Associated microRNAs and the Cholinergic Anti-inflammatory Pathway upon LPS-Treatment in Rats. J Neuroimmune Pharmacol, (2022). 17(1-2): p. 261–276. DOI: 10.1007/s11481-021-10003-w.

40. van Norden J, Spies CD, Borchers F, Mertens M, Kurth J, Heidgen J, Pohrt A, Mueller A, The effect of peri-operative dexmedetomidine on the incidence of postoperative delirium in cardiac and non-cardiac surgical patients: a randomised, double-blind placebo-controlled trial. Anaesthesia, (2021). 76(10): p. 1342–1351. DOI: 10.1111/anae.15469.

41. Erecinska M Dagani F, Relationships between the neuronal sodium/potassium pump and energy metabolism. Effects of K+, Na+, and adenosine triphosphate in isolated brain synaptosomes. J Gen Physiol, (1990). 95(4): p. 591–616. DOI: 10.1085/jgp.95.4.591.

42. Ivannikov MV, Sugimori M, Llinas RR, Calcium clearance and its energy requirements in cerebellar neurons. Cell Calcium, (2010). 47(6): p. 507–13. DOI: 10.1016/j.ceca.2010.04.004.

43. Silver IA Erecinska M, Energetic demands of the Na+/K+ ATPase in mammalian astrocytes. Glia, (1997). 21(1): p. 35–45. DOI: 10.1002/(sici)1098-1136(199709)21:1<35::aid-glia4>3.0.co;2-0.

